# Dimerization processes for light-regulated transcription factor Photozipper visualized by high-speed atomic force microscopy

**DOI:** 10.1101/2022.03.27.485139

**Authors:** Akihiro Tsuji, Hayato Yamashita, Osamu Hisatomi, Masayuki Abe

## Abstract

Dimerization is critical for transcription factors (TFs) to bind DNA and regulate a wide variety of cellular functions; however, the single-molecular mechanisms remain to be completely elucidated. Here, we used high-speed atomic force microscopy (HS-AFM) to observe the dimerization process for a photoresponsive TF Photozipper (PZ), which consists of light-oxygen-voltage-sensing (LOV) and basic-region-leucine-zipper (bZIP) domain. HS-AFM visualized not only the oligomeric states of PZ molecules forming monomers and dimers under controlled dark–light conditions but also the domain structures within each molecule. Successive AFM movies captured the dimerization process for an individual PZ molecule and the monomer–dimer reversible transition during dark–light cycling. The high-resolution AFM images of the domain structures in PZ molecules demonstrated that the bZIP domain entangled under dark conditions was loosened owing to light illumination and fluctuated around the LOV domain. These observations revealed the role of the bZIP domain in the dimerization processes of a TF.

## Introduction

Transcription factors (TFs) are essential DNA binding proteins that regulate a wide range of gene expression. Today, as many as ~1,600 TFs in the human genome^1^ and ~48,000 TFs in the eukaryote genome^2^ have been identified. They control various cellular functions critical for living organisms, such as cell proliferation, homeostasis, and immune response^3,4^. A eukaryotic TF monomer recognizes short DNA sequences of approximately 6–8 nucleotide pairs. In addition, these short sequences can randomly occur many times in large genomes. Such numerous random occurrences can lead to spurious binding of TFs^5^. To express the right gene at the right location, at the right moment and at the right level, high specificity must be present for the target DNA sequences. Many TFs form dimers to achieve such high specificity^4,6^. This dimer formation doubles the total effective length of the target sequence and greatly increases both the affinity and the recognition specificity of TF binding compared with those of a monomer^7^. For example, the binding affinity of Gal4, a transcriptional activator in a yeast, for the target DNA has been reported to be dramatically reduced by deletion of the dimerization domain^8,9^. Thus, elucidating the TF dimerization mechanism is critical for understanding not only the structure–function relationship of TFs but also the general principle of transcription.

TFs are often categorized into families depending on their dimerization domain type. The basic-region-leucine-zipper (bZIP) domain is one type, and forms a major family^5,7^. Many TFs contain an additional dimerization domain conferring specificity in the range of interactions, such as Per–ARNT–Sim (PAS) domains for signal sensing^4,10,11^. One typical PAS domain is the light–oxygen–voltage-sensing (LOV) domain, which has been found in numerous species and extensively studied^12^. Aureochrome-1 (AUREO1) is a TF possessing both the bZIP domain and the LOV domain, and it has been reported to play roles in blue light (BL)-induced branching and cell proliferation in a stramenopile alga, *Vaucheria frigida*^13–15^. In addition, AUREO1 has been found to dimerize under BL illumination and to bind to the TGACGT target DNA sequence, indicating that this protein is a BL-responsive TF^13,16^. Photozipper (PZ), an N terminally truncated and modified AUREO1, has been constructed for development as an optogenetic tool, and the BL-induced dimerization mechanism has been investigated^16–18^. Size-exclusion chromatography (SEC) and the electrophoretic mobility shift assay revealed that PZ forms a dimer under BL illumination and increases its affinity for the target DNA sequence^16,19^. By studying bZIP deletion mutant using dynamic light scattering (DLS) and fluorescence resonance energy transfer (FRET), researchers have proposed a model for BL-induced dimerization of PZ^17^. According to the model, the bZIP domain shields the dimerization site of the LOV domain under dark conditions, and the LOV–bZIP intramolecular interaction stabilizes the monomeric form. By contrast, the BL-induced conformational changes in the LOV domain weaken the LOV–bZIP interaction and cause the dissociation of the bZIP domain from the LOV domain. PZ subsequently dimerizes via bZIP–bZIP and LOV–LOV intermolecular interactions^17^. A structural analysis of only the LOV domain in an aureochrome from *Phaeodactylum tricornutum* has clarified the BL-induced conformational changes within the LOV domain^20^, and the site-directed mutant studies of PZ have confirmed that the conformational change can affect the LOV–bZIP interaction^18,21^. These studies support some parts of the previously described model. However, in a series of dimerization processes, whole structural arrangement of a PZ molecule remains unrevealed in each state because aureochromes, including PZ, have thus far eluded structural elucidation at full length because of the existence of disordered regions^20,22–24^. In particular, the structural configuration of the bZIP domain in a PZ molecule is speculated to play a key role in the dimer formation^17^. Therefore, a study of the single-molecular structure and dynamics for PZ at full length is necessary.

High-speed atomic force microscopy (HS-AFM) has been extensively used to visualize various biological molecules in physiological solution at nanoscale resolution^25–27^. Notably, HS-AFM observations of intrinsically disordered proteins have revealed flexible and fluctuating unstructured regions^28–30^. These studies have demonstrated that HS-AFM is a useful technique for the single-molecule-level analysis of such proteins containing unstructured segments. Recently, GAL4-VVD, a chimeric fusion construct of a photoresponsive TF^31^, has been observed on DNA origami by HS-AFM^32^. Such studies have focused on the dynamics of the fusion protein during the search for the target sites on DNA. However, detailed observations of the TF dimerization process that play an important role in DNA binding have not been achieved.

Here, we carried out high-resolution observations of the photoresponsive TF “Photozipper” using HS-AFM to elucidate the dimerization mechanism. The HS-AFM images clearly visualized not only the oligomeric states of PZ molecules forming monomers and dimers but also the domain structures within each molecule. Successive AFM movies captured the dimerization process for an individual PZ molecule and the monomer–dimer reversible transition on light–dark cycles. We found that the dimer form is very stable under light conditions compared with the form under dark conditions. The AFM analysis of the domain structures in PZ molecules demonstrated that the bZIP–linker region entangled under dark conditions is loosened because of light illumination. Our results highlight the role of the bZIP domain on PZ dimerization through BL illumination. In addition, they advance understanding of the dimerization mechanism in a TF at the single-molecular level.

## Results

### Preparation of recombinant PZ proteins

We prepared a recombinant PZ protein by substituting serine for Cys162 and Cys182 in an N-terminally truncated AUREO1 of *Vaucheria frigida* (Fig. 1a). We also prepared a mutant protein PZ-S_2_C by substituting cysteine for Ser162 and Ser182 in wild-type PZ^33^ (Fig. 1b). Mutant S_2_C can stably dimerize by forming intermolecular disulfide linkages with the two cysteines. This mutant was used for comparing the molecular structure and volume with dimeric PZ. Fig. 1c shows the sodium dodecyl sulfate polyacrylamide gel electrophoresis (SDS-PAGE) results for the wild-type PZ and mutant S_2_C samples. The results for each sample show a single band corresponding to a molecular weight of ~30 kDa for PZ and ~60 kDa for S_2_C. These results indicate that each sample was sufficiently pure for use in single-molecule imaging by HS-AFM and that mutant S_2_C stably formed the dimer.

**Figure 1:**
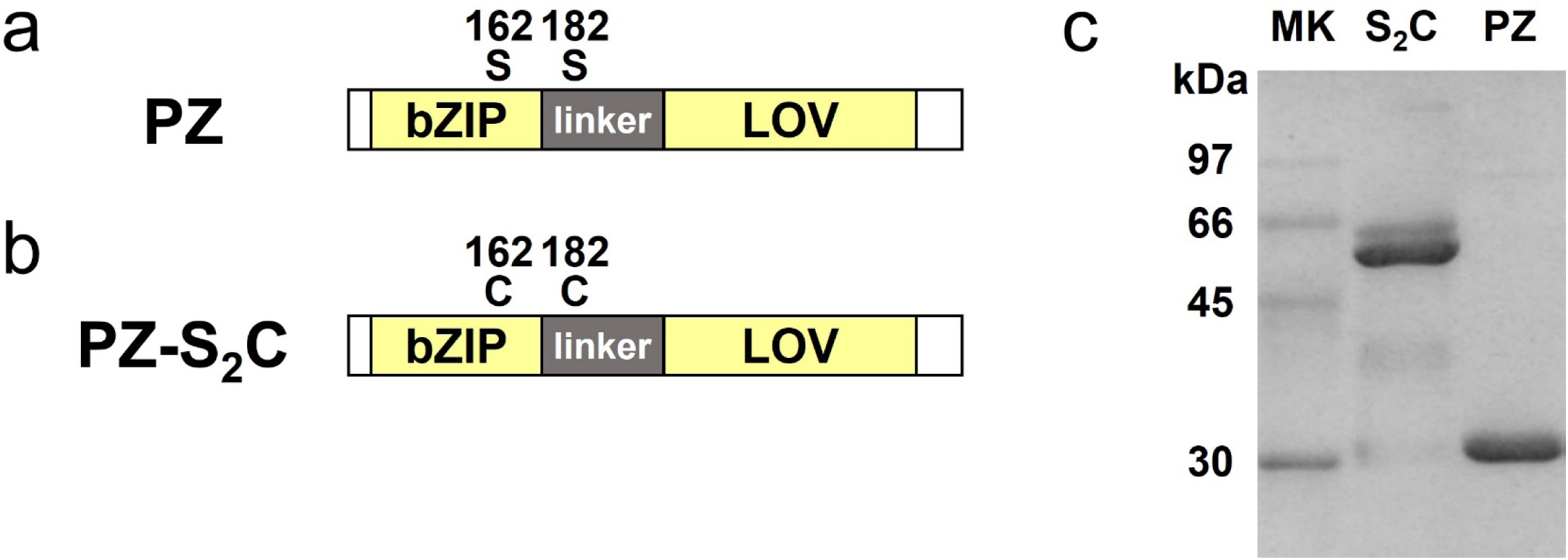
Schematic diagrams and SDS-PAGE of recombinant PZ proteins. (a, b) Schematic diagrams of the domain structure in PZ and PZ-S_2_C. Two serine residues in PZ and their substituted cysteine residues in PZ-S_2_C are shown with sequence number. (c) SDS-PAGE of recombinant proteins: Marker (left lane), PZ-S_2_C (center lane), PZ (right lane).

### HS-AFM observations of PZ molecules

We observed the purified PZ molecules using HS-AFM in physiological solution under dark and light conditions. Fig. 2a shows a still image from a successive HS-AFM movie for wild-type PZ captured under dark conditions (Supplementary Movie 1). Globular-shaped molecules ~5 nm in diameter were visualized in the AFM image, and they were extensively diffused on the mica surface (see also Supplementary Movie 1). Occasionally, molecules ~10 nm in diameter were also observed (see Supplementary Fig. 1). Fig. 2b shows a still image of the HS-AFM movie for wild-type PZ captured under continuous BL illumination (Supplementary Movie 2). Globular-shaped molecules with two diameters were observed in the AFM image. The smaller ones were ~5 nm in diameter, and the larger ones were ~10 nm. The smaller ones rapidly diffused on the mica surface, whereas the larger ones slowly diffused (see also Supplementary Movie 2). In this HS-AFM movie, a tail-like structure was observed to fluctuate around a globular structure in the larger molecules. To compare these molecular structures with the stable PZ dimer, we observed dimeric mutant protein S_2_C using HS-AFM. Fig. 2c shows a HS-AFM image of the mutant S_2_C molecules. The globular molecules were observed, and they exhibited similar diameters to the larger molecules of wild-type PZ under illumination. These molecules slowly diffused on the mica surface (Supplementary Movie 3). In contrast to wild-type PZ, the smaller molecules were not observed in the HS-AFM movie of mutant S_2_C. To quantitatively compare these results, we analyzed the volumes of the molecules in the HS-AFM images. Fig. 2d–f shows histograms for the molecular volumes analyzed from the HS-AFM images of wild-type PZ under dark and light conditions and from the images of mutant S_2_C. The histograms of wild-type PZ under dark conditions (Fig. 2d) and light conditions (Fig. 2e) show bimodal distributions, whereas that of mutant S_2_C shows a unimodal distribution (Fig. 2f). The mean volume of the mutant S_2_C was in good agreement with the major distribution of higher volumes of wild-type PZ under light conditions. In addition, the major distribution of wild-type PZ under dark conditions was in good agreement with the minor distribution of lower volumes of wild-type PZ under light conditions. Therefore, we conclude that the major molecules observed under dark conditions are PZ monomers and that the larger molecules observed under light conditions are PZ dimers. Here, we note that the minor distribution of the larger molecules is seen in Fig. 2d. These molecules are dimers under dark conditions. The results of volume analyses also show that PZ predominantly exists as a monomer under dark conditions and as a dimer under light conditions. These results are consistent with a previous study involving SEC^17^, which revealed oligomeric states of PZ under each condition. Thus, we demonstrated that PZ monomers and dimers can be distinguished by HS-AFM.

**Figure 2:**
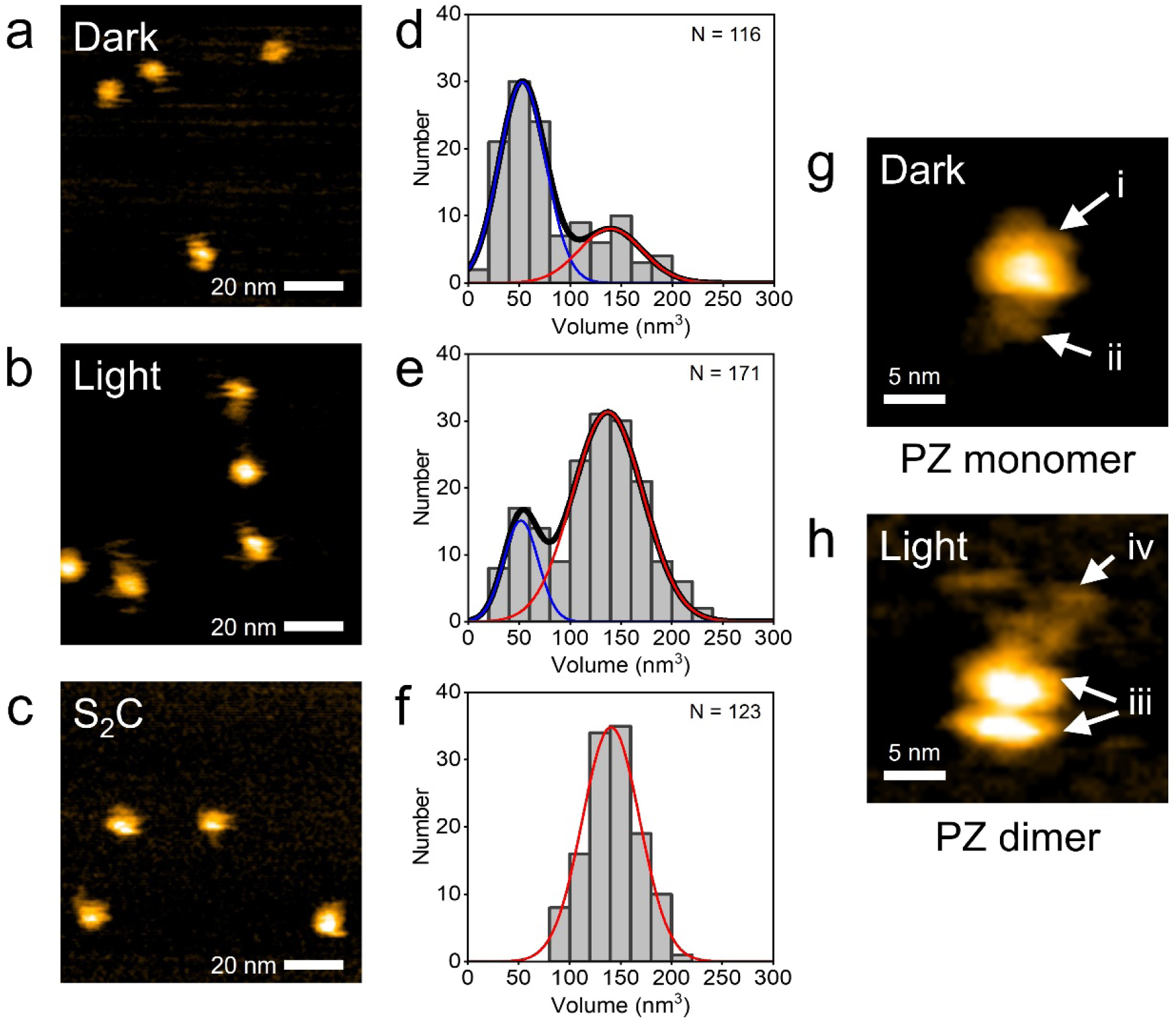
HS-AFM observations of PZ molecules. (a, b) HS-AFM images of PZ molecules observed under dark and light conditions, respectively. (c) HS-AFM image of mutant protein PZ-S_2_C observed under light conditions. These show still images of successive AFM movie. See also Supplementary Movies 1,2 and 3. Several PZ and PZ-S_2_C molecules were observed on mica surface. (d-f) Histograms for the molecular volumes analyzed from AFM images of wild-type PZ under dark, light condition and S_2_C mutant. Curve represents the fit to a Gaussian distribution. Mean value of each Gaussian curve: (d) 1st peak = 52.9 nm^3^, 2nd peak = 139 nm^3^, (e) 1st peak = 52.4 nm^3^, 2nd peak = 137 nm^3^, (f) 141 nm^3^. (g) Highly magnified AFM image of PZ monomer under dark conditions. This molecule was included in the major distribution in Fig. 2d. The folded-string structure was seen around the spherical structure. (h) Highly magnified AFM image of PZ dimer under light conditions. This molecule was included in the major distribution in Fig. 2e. The dimer exhibited a bilobed structure together with the tail-like structure. Protein concentration: (a) 25 nM, (b) 50 nM, (c) 50 nM. Scan ranges: (a, b and c) 100×100 nm^2^, (g and h) 24.5×24.5 nm^2^. Scan rates: (a and g) 1 sec/frame, (b, c and h) 0.5 sec/frame.

We next focused on the structures of the PZ molecules. Fig. 2g shows a magnified HS-AFM image of a PZ monomer under dark conditions. This molecule is included in the major distribution in Fig. 2d. Two discernible structural domains of the PZ molecule are observed in the HS-AFM image: a “spherical structure” and a “folded string-like structure,” as indicated by labels “i” and “ii” in Fig. 2g, respectively. As shown in Fig. 1a, PZ is composed of two main structural domains: LOV and bZIP, which are connected by a linker region. Previous crystal-structure analyses have revealed that the LOV domain in AUREO1 is spherical, consisting of α-helices and β-sheets^20,22^. Therefore, the spherical structure in our AFM image is thought to be the LOV domain. On the other hand, the folded string-like structure in the AFM image turns out to be both the bZIP domain and the linker region (bZIP-linker region). Analyses by circular dichroism (CD) spectroscopy and nuclear magnetic resonance (NMR) spectroscopy have shown that the bZIP domain partially forms α-helices in a DNA unbound state^19,34,35^. However, the other structures except for the α-helices remain unrevealed in the bZIP-linker region, although numerous structural studies have been reported^20^. This situation implies that their unknown structures could be disordered in the DNA unbound state^22,24^. In the present study, we visualized the total architecture of the bZIP-linker region under dark conditions for the first time.

Fig. 2h shows a magnified HS-AFM image of a PZ dimer under illumination, which is included in the major distribution in Fig. 2e. Two discernible structural domains of the PZ molecule are evident in the HS-AFM image: a “bilobed structure” and a “fluctuated tail-like structure,” as indicated by labels “iii” and “iv” in Fig. 2h, respectively. Previous SEC studies of isolated LOV domains have suggested that the LOV domain dimerizes under illumination.^17,36^ Also, a previous crystal-structural analysis revealed that two LOV domains pair as a dimer under illumination^20^. Therefore, the bilobed structure in our AFM image is thought to be the dimerized LOV domain. The size of the bilobed structure is actually larger than that of the LOV domain in PZ monomers. Here, we note that such a dimerized structure of the LOV domain fluctuates between bilobed structures and globular structures (see Supplementary Fig. 2 and Supplementary Movie 4). This observation suggests that the LOV–LOV interaction varies with time. On the other hand, it turns out that the fluctuated tail-like structure in the AFM image is the bZIP-linker region. In a previous study, SEC and FRET analyses revealed that bZIP domains of wild-type PZ also dimerize under illumination^17^. Although we could not resolve bZIP domains dimerizing in our HS-AFM observations, our AFM movies revealed for the first time that they were unexpected flexible structures that fluctuate around the LOV domain (Supplementary Movie 2).

### Dynamic movement analysis of PZ molecules

The HS-AFM movies showed obvious differences in not only the molecular size but also the dynamic behavior of PZ molecules under dark and light conditions. PZ monomers rapidly diffused on a mica surface under dark conditions (Supplementary Movie 5). By contrast, PZ dimers fluctuated, exhibiting slow diffusion under illumination (Supplementary Movie 6). To compare these dynamic behaviors, we analyzed the trajectories of PZ molecules in AFM movies. Representative trajectories of PZ monomers under dark conditions and PZ dimers under illumination are shown in Fig. 3a and b, respectively. These PZ molecules exhibited random movements, indicating that the movements are not the effect of a tip– sample interaction force but rather molecular diffusion on the mica surface in a buffer solution. The diffusion speeds of the PZ molecules were quantified by calculating the mean-square displacements (MSDs) for each trajectory. As shown in Fig. 3c and d, each MSD linearly increases with time. In our HS-AFM observations, PZ molecules diffused in two dimensions on the mica surface. Therefore, the MSD plots can be fitted with the following equation for two-dimensional random diffusion^37^:

**Figure 3:**
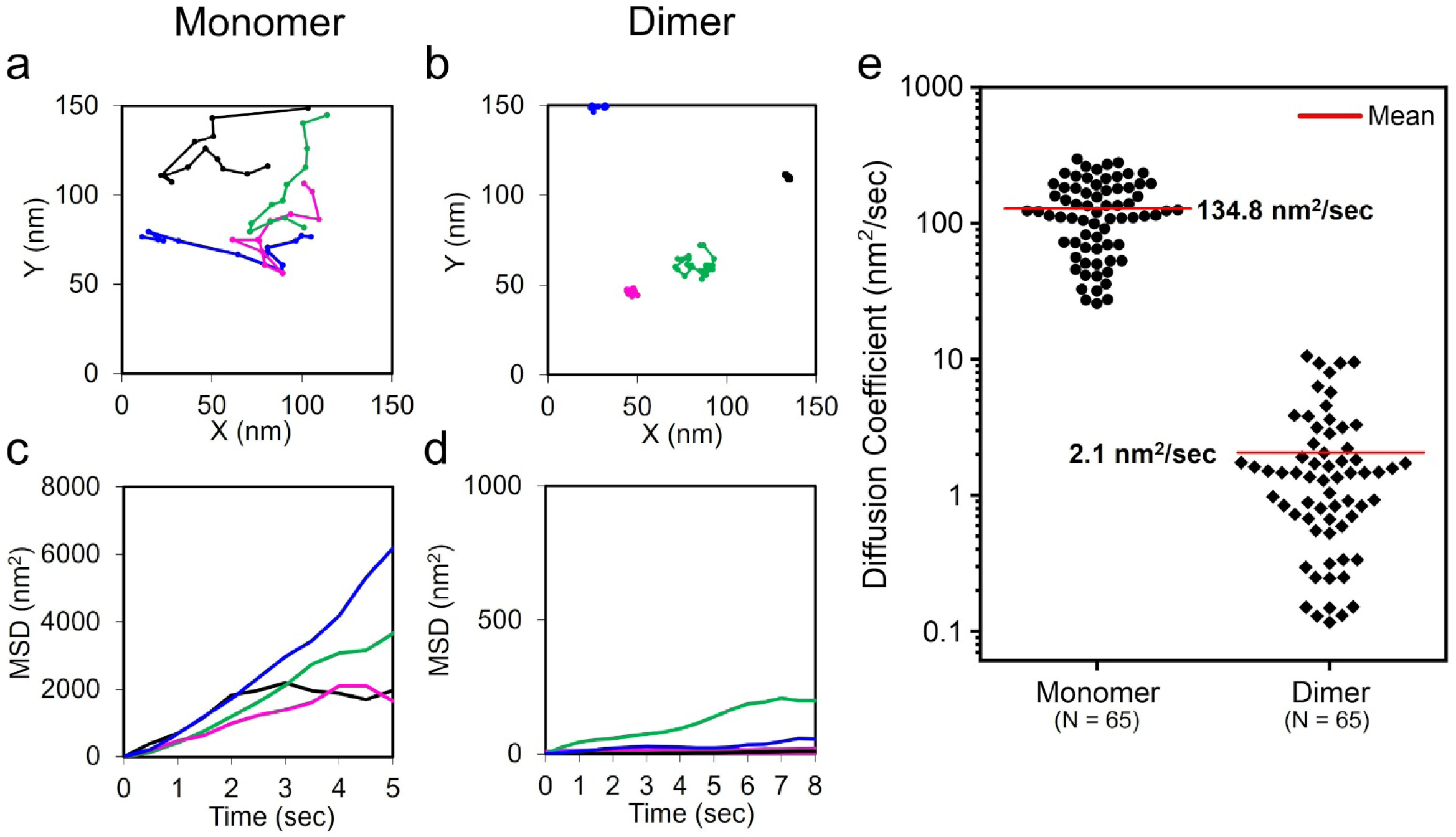
Diffusion analysis of PZ monomer and dimer. (a) Molecular trajectories of the two-dimensional diffusion for PZ monomers on mica surface under dark conditions. Four PZ monomers are shown in this graph, which were chosen from the major distribution in Fig.2d. The mass center position of each PZ molecule was analyzed and tracked in Supplementary Movie 5. (b) Molecular trajectories of the two-dimensional diffusion for PZ dimers under light conditions. Four PZ dimers are shown in this graph, which were chosen from the major distribution in Fig.2e. The mass center position of each PZ molecule was analyzed and tracked in Supplementary Movie 6. (c, d) Mean square displacements (MSD) analyzed from individual trajectories of (a) and (b), respectively. These MSD linearly increased with time, yielding diffusion constants. (e) Beeswarm plots of diffusion constants analyzed from individual trajectories. Mean value of each distribution: (Monomer) 134.8 nm^2^/sec, (Dimer) 2.1 nm^2^/sec. Protein concentration: (a, b) 13nM.

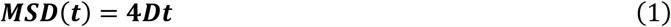

where t is time and *D* is the diffusion coefficient. The diffusion coefficients for individual PZ molecules are shown in Fig. 3e. The average diffusion coefficient of the dimer was approximately 70 times lower than that of the monomer. This work represents the first observation of the dynamic behavior of each PZ molecule under dark and light conditions using HS-AFM. In a previous study based on DLS, the hydrodynamic radii (*R*_H_) of PZ molecules were observed to increase under illumination^17^. Here, higher *R*_H_ values correspond to lower diffusion constants from the Stokes–Einstein relationship^38^. Therefore, this result suggests that the diffusion speeds of PZ molecules decreased upon light-induced dimerization. Our result is consistent with the results of the previous study showing that the diffusion coefficient of the PZ dimer is lower than that of the PZ monomer^17^. It should be noted that the two-dimensional mobility of PZ molecules on the mica surface is much lower than the three-dimensional mobility estimated from previous biophysics experimental data^17,39^. The interaction of PZ molecules with the mica substrate can affect the diffusion speed. Moreover, the average diffusion coefficient of the PZ monomers differed significantly from that of the dimers in the present study. Although we could not elucidate a major cause of the large difference, the conformational change due to the dimer formation of PZ molecules may be related to the decrease of the diffusion speed. We also note that an aligned orientation of LOV domains in the dimer form can affect the interaction with the mica substrate.

### Photoinduced transition process for PZ molecules from monomer to dimer

To investigate the dynamic transition process from PZ monomers to dimers, we conducted successive HS-AFM observations of PZ molecules as the conditions were changed from dark to light. Fig. 4a shows snapshots of a successive HS-AFM movie (Supplementary Movie 7). Before BL illumination, PZ monomers were predominant and highly diffusing on the mica surface (dark in Fig. 4a). After BL illumination, the number of PZ dimers increased with time (HS-AFM images from 140 sec to 342 sec in Fig. 4a) and the diffusion speed on the mica surface decreased upon dimer formation (see also Supplementary Movie 7). Finally, PZ dimers became predominant in the AFM image at 390 sec. In these successive AFM measurements, we observed the transition processes for PZ molecules from monomers to dimers. Fig. 4b shows snapshots of high-magnification AFM images captured during the dimerization processes under BL illumination. Two PZ monomers were diffusing on the mica surface, as shown in the AFM images at 0 sec and 2.5 sec in Fig 4b. These monomers collided with each other and formed a single dimer in the AFM image at 4 sec. After the molecular collision, the PZ dimer remained bound at 7.5 sec (see also Supplementary Movie 8).

**Figure 4:**
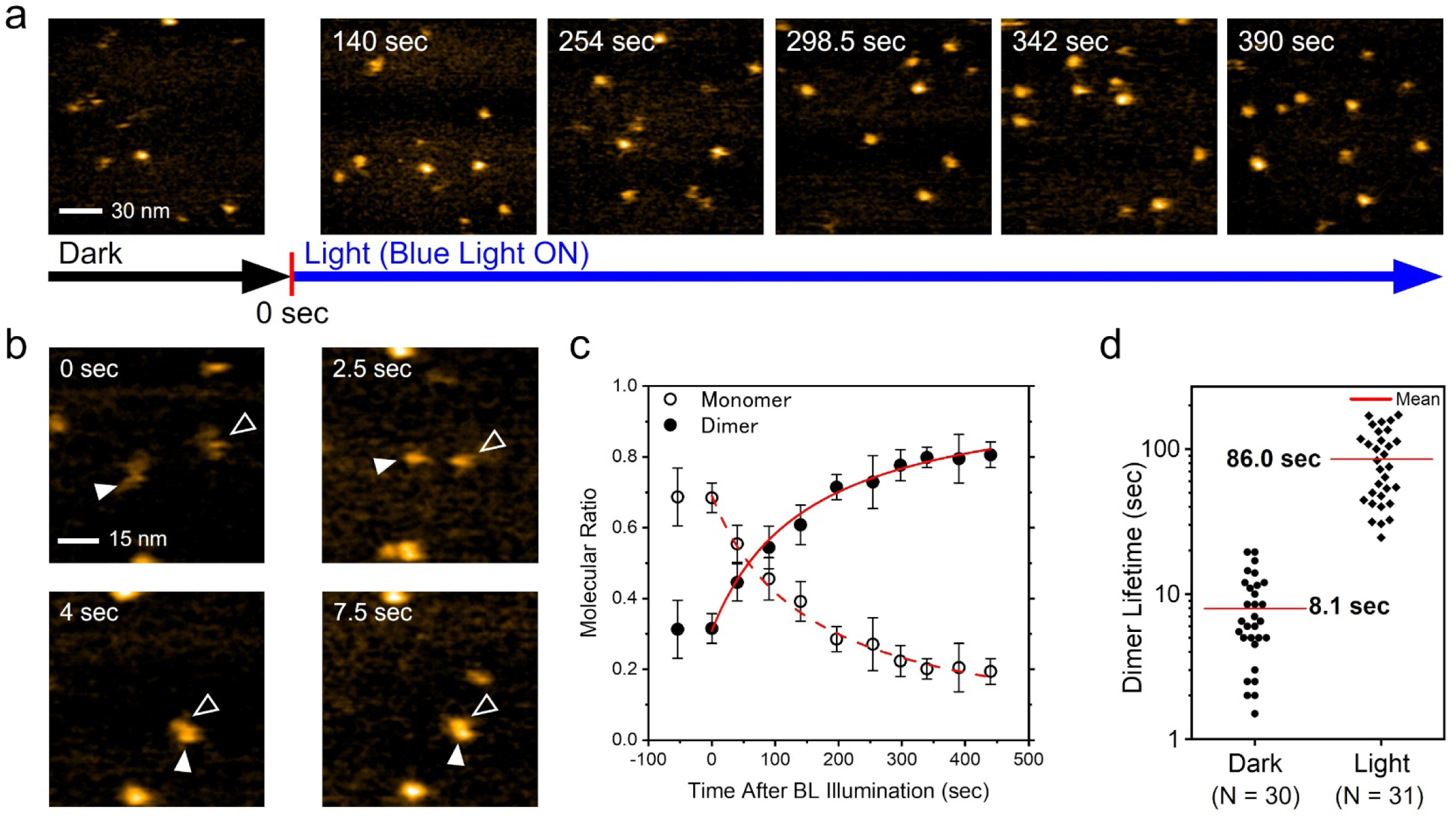
Photoinduced transition process of PZ molecules from monomer to dimer observed by HS-AFM. (a) HS-AFM observation of photoinduced transition process of PZ molecules from dark to light state. Time displays in AFM images indicate time course after blue light (BL) illumination. The number of PZ dimers increased with time (see also Supplementary Movie 7). (b) Snapshots of successive high magnification AFM movie captured dimerization processes of PZ molecules under BL illumination (Supplementary Movie 8). Two PZ monomers were diffusing on mica surface at 0 sec and 2.5 sec. Each open and filled arrowheads indicate individual PZ monomers. At 4 sec, two PZ monomers collided with each other and formed one dimer. The dimer remained bound at 7.5 sec. (c) Time courses of each ratio of monomer and dimer detected in AFM images. Each ratio was normalized by the total PZ molecules in each AFM image. Note that one PZ dimer was counted as two PZ molecules. Open and filled circles indicate average ratio of monomer and dimer, respectively (N = 3). Error bars indicate the standard deviation. These plots were fitted with equations (12) and (13) in the Supplementary Note, respectively (red solid line and dotted line). (d) Beeswarm plots of dimer lifetimes of PZ molecules under dark and light conditions. These data were analyzed from successive AFM measurement before and after light illumination. Mean value of each distribution: (Dark) 8.1 seconds, (Light) 86.0 seconds. Protein concentration: (a and b) 38 nM. Scan ranges: (a) 150×150 nm^2^, (b) 75×75 nm^2^. Scan rates: (a and b) 0.5 sec/frame.

We analyzed the number of monomers and dimers at each time point in the successive AFM movies to investigate the kinetics of such a transient response to illumination. Fig. 4c shows time courses of each ratio of monomers and dimers among total molecules. Under dark conditions before BL illumination, the monomer ratio was predominant. Moreover, each ratio of monomers and dimers remained constant at different time points. This result indicates that the monomer and the dimer were in equilibrium. After BL illumination, the dimer ratio gradually increased with time, whereas the monomer ratio decreased with time. The dimer ratio exceeded the monomer ratio at ~60 sec. Finally, each ratio became saturated, and the dimer ratio was predominant under continuous-light conditions. These results indicate that the equilibrium of PZ transitioned from the monomer-predominant dark state to the dimer-predominant light state because of BL illumination (see also Supplementary Note). The transient change of monomers and dimers in such an equilibrium shift can be described by equations (12) and (13), respectively, in the Supplementary Note. Our data were well fitted with these equations, as shown by solid and dashed curves in Fig. 4c. The exponential time constant (τ) was 255 ± 26 sec, as calculated using equation (14) in the Supplementary Note. A previous study that used a transient grating (TG) method revealed that the time constant of PZ dimerization was 2 sec at a PZ concentration of 20 μM^39^. This concentration is three orders of magnitude higher than ours (38 nM). The PZ dimerization speed depends on the PZ concentration^39^: a high concentration leads to a low time constant (see equation (14) in the Supplementary Note). Therefore, our low PZ concentration is considered to result in an increase of the time constant. The TF concentrations in cells are known to be on the order of 10 to 100 nM^40,41^. We successfully measured PZ behavior by HS-AFM under concentration conditions similar to the concentrations in cells. It should be noted that the diffusion speed of PZ molecules are decreased in our measurements owing to the interaction between the PZ and the mica substrate. Here, we performed successive HS-AFM measurements of PZ kinetics in repeated light–dark cycles to confirm the reproducibility of the photoinduced transition between the monomers and dimers (Supplementary Fig. 3). The initial BL responses were approximately the same as that in Fig. 4c. Under dark conditions, after the initial BL illumination, the dimer ratio gradually decreased with time, whereas the monomer ratio increased with time. At ~1300 sec, each PZ ratio returned to the initial state. Similar responses were reproduced in the second light–dark cycle. Therefore, these results indicate that such a photoinduced monomer–dimer transition of PZ is a reversible process. We demonstrated that HS-AFM can reproducibly capture the physiological behavior of PZ at the single-molecule level.

Successive HS-AFM movies captured not only the processes of dimer formation but also that of dissociation (see Supplementary Fig. 4 and Supplementary Movie 9). Through these measurements of the photoinduced transition processes, we have noticed an interesting difference in that the PZ dimer formation is very stable under BL illumination compared with that under dark conditions. We therefore analyzed the lifetimes from dimer formation to dissociation. Fig. 4d shows the dimer lifetimes under dark and light conditions. Notably, these dark dimers are not in a transient state but in an equilibrium state under dark conditions. The average dimer lifetime under illumination (86.0 sec) was 10 times longer than that under dark conditions (8.1 sec). This result suggests that the intramolecular interaction of the PZ dimer (monomer–monomer intermolecular interaction) under BL illumination is much stronger than that under dark conditions. This work is the first report directly comparing the stability of each PZ dimer molecule under dark and light conditions.

### Light-induced conformational change of PZ monomers

We found that the PZ dimer is more stable under BL illumination than under dark conditions. This increase in dimer stability is attributed to the conformational change of PZ molecules under illumination. To reveal the mechanism of dimer stabilization, we focused on the molecular structure of the PZ monomer under light and dark conditions. Fig. 5a shows a magnified HS-AFM image visualizing the high-resolution structure of a PZ monomer under dark conditions. As described in Fig. 2g, the spherical structure corresponds to the LOV domain and the folded string-like structure corresponds to the bZIP-linker region. Fig. 5b shows a HS-AFM image gallery of the PZ monomers under dark conditions. The structure of each monomer is similar to that in Fig. 5a, and such structures were also confirmed for the other monomers under dark conditions. Fig. 5d shows a magnified HS-AFM image of a PZ monomer under illumination. Surprisingly, an unfolded string-like structure anchored to the spherical LOV domain is visualized in this high-resolution HS-AFM image. Fig. 5e shows a HS-AFM image gallery of PZ monomers under illumination. The structure of each monomer was similar to that in Fig. 5d, and these structures appeared to wobble on the mica surface. Such structures were also confirmed for the other monomers under light conditions. These results suggest that the illumination changed the conformation of the PZ monomers.

**Figure 5:**
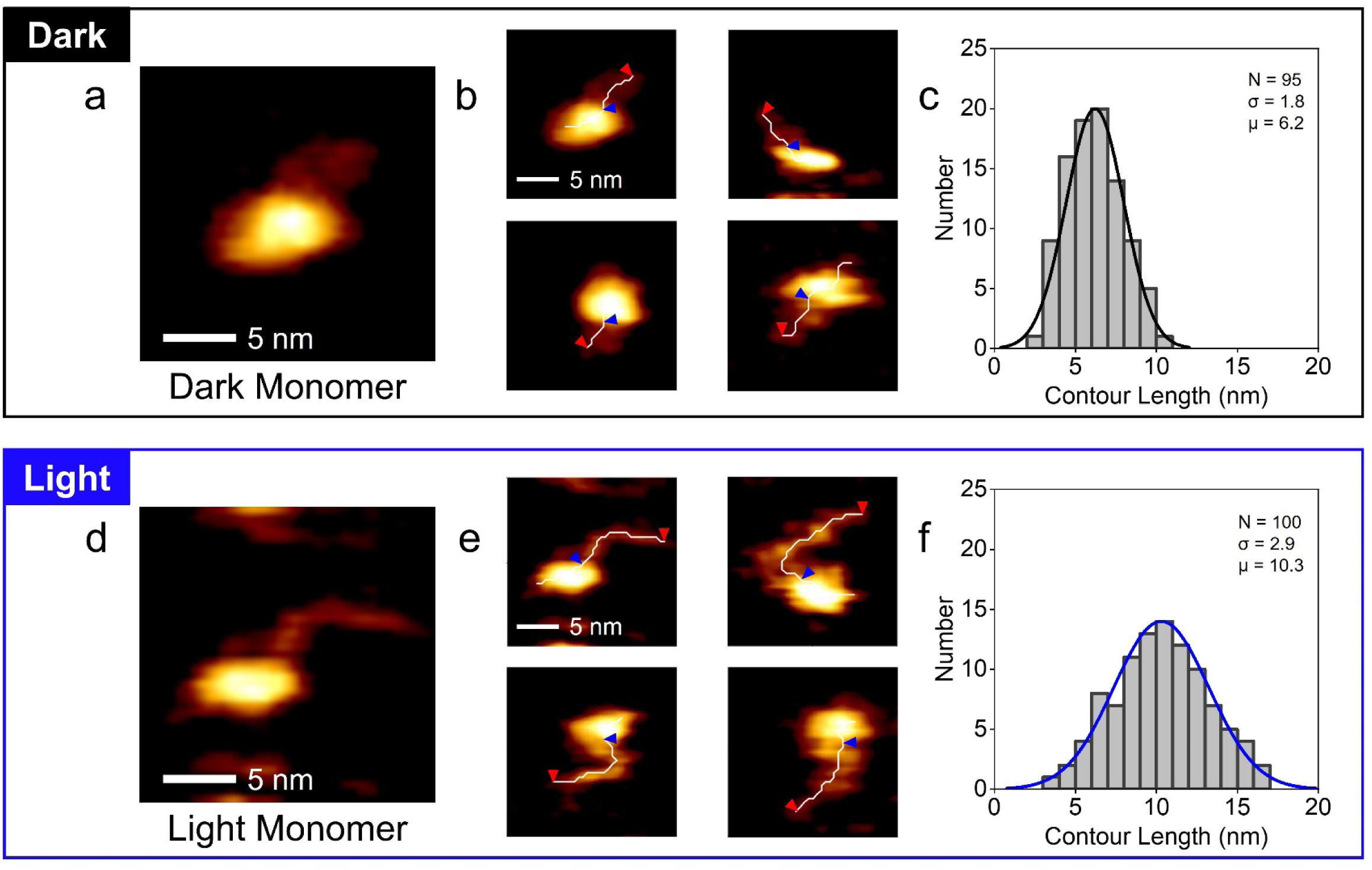
HS-AFM observations of PZ molecules under dark and light conditions. (a) Typical HS-AFM image of PZ monomer under dark conditions. (b) HS-AFM image gallery of PZ monomers under dark conditions. A skeleton line is drawn in white on each PZ monomer. Red and blue arrowheads show each end of the bZIP-linker region in PZ monomer (see Methods and Supplementary Fig. 5). The lengths of the skeleton lines between red and blue arrowheads were analyzed as the contour lengths of the bZIP-linker region. (c) Histograms for the bZIP-linker contour lengths analyzed from AFM images of PZ monomers under dark conditions. Curve represents the fit to a Gaussian distribution. The mean value of the Gaussian curve: μ = 6.2. (d) Typical HS-AFM image of PZ monomer under light conditions. (e) HS-AFM image gallery of PZ monomers under light conditions. The skeleton lines and the contour lengths were analyzed by same procedures as (b). (f) Histograms for the bZIP-linker contour lengths analyzed from AFM images of PZ monomers under light conditions. The mean value of the Gaussian curve: μ = 10.3. Protein concentrations: (a, b and c) 13 or 25 nM, (d, e and f) 25 or 50 nM. Scan ranges: 20×20 nm^2^, Scan rates: 0.5 sec/frame.

To investigate the conformational change of the PZ, we analyzed the contour lengths of string-like structures in the PZ monomers, as shown by white lines and two arrowheads in Fig. 5b and e (see also Methods and Supplementary Fig. 5). Fig. 5c and f show the histograms for the contour lengths analyzed from the HS-AFM images of the PZ monomers under dark and light conditions, respectively. The mean value of the contour lengths under illumination (10.3 nm) was 1.7 times longer than that under dark conditions (6.2 nm). In addition, the distribution of the contour lengths under illumination (*σ* = 2.9) was wider than that under dark conditions (*σ* = 1.8). These results indicate that the string-like structure that was folded under dark conditions was unfolded under illumination and then fluctuated in solution. Thus, we conclude that the bZIP-linker region entangled under dark conditions is loosened under illumination.

## Discussion

We observed the single-molecular structure and dynamics of PZ in physiological buffer solution using HS-AFM under controlled dark–light conditions. The HS-AFM images show monomer dominance under dark conditions and dimer dominance under light conditions. HS-AFM movies captured the dimerization process for individual PZ molecules and the monomer–dimer reversible transition during dark–light cycles. For the first time, high-resolution HS-AFM was used to visualize detailed structures of PZ and revealed the structural configuration of the bZIP domain in the full-length PZ molecule in both the light state and the dark state. These results demonstrate the previous PZ dimerization model^17^ and update details of some aspects of the model, as described in Fig. 6.

**Figure 6:**
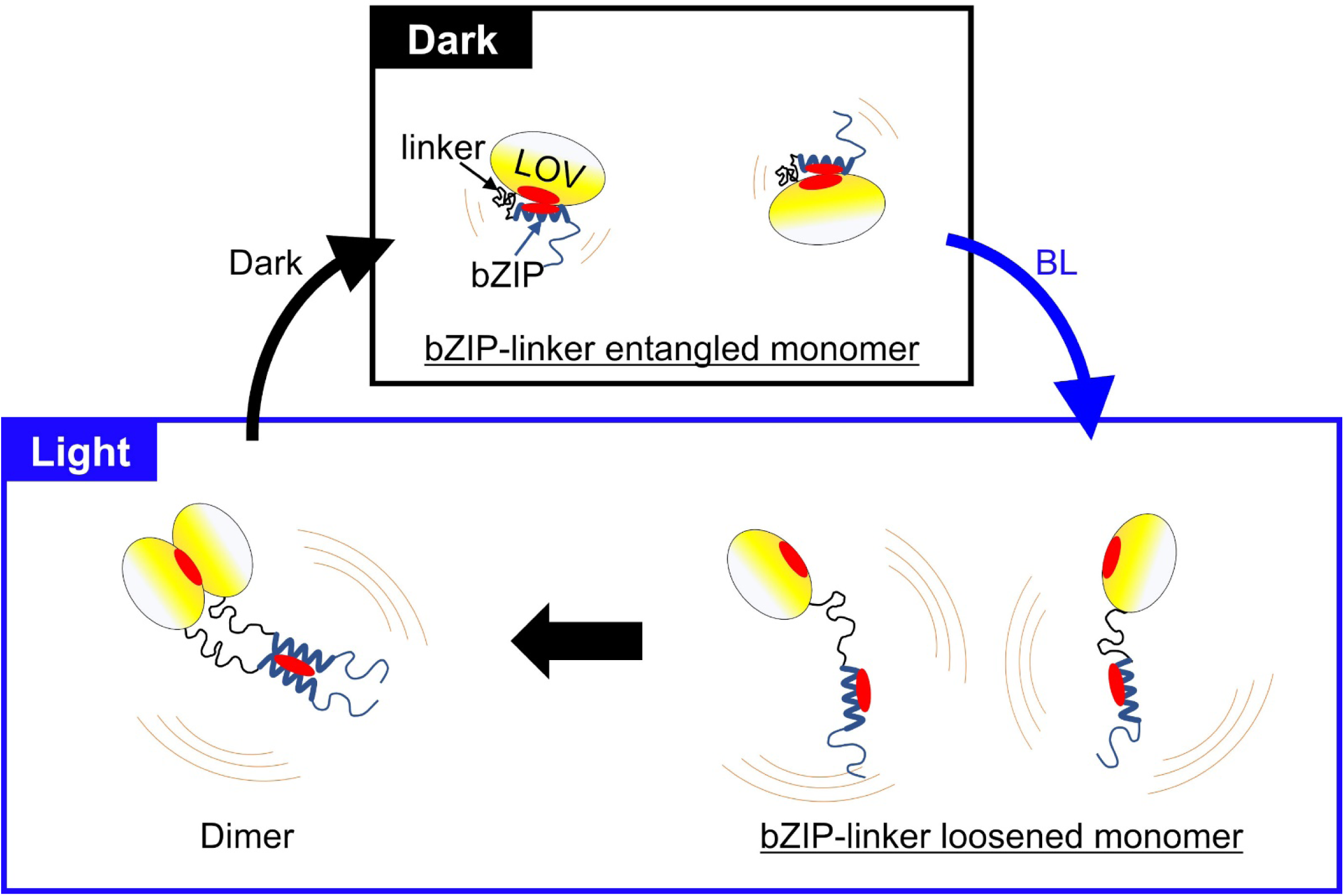
Schematic model of the BL induced dimerization process of Photozipper. Under dark conditions, PZ exists as a bZIP-linker entangled monomer where the hydrophobic dimerization sites are hidden by the bZIP–LOV intramolecular interaction. Such molecular configuration of PZ inhibits the dimerization under dark conditions. BL induces the dissociation of the bZIP from the LOV surface and the dimerization sites become exposed. The loosened and elongated bZIP-linker fluctuates around the LOV. This flexible and dynamic movement of bZIP increases the contact with the other monomers and facilitates the dimerization. The intermolecular interaction of LOV–LOV and bZIP–bZIP stabilize the PZ dimer.

In the dark state, the bZIP-linker region is entangled and interacts with the LOV domain. Here, we considered that the bZIP domain binds to the LOV domain, hiding the dimerization sites of both the LOV domain and the bZIP domain, as previously reported^20^. Therefore, the dimerization with the other PZ monomer is inhibited under dark conditions. Indeed, we found that the binding lifetime of the formed dimer was very short under dark conditions compared with that under illumination (Fig. 4d). Thus, the bZIP domain and the LOV domain play a role in mutual autoinhibition of PZ dimerization in the dark state. By contrast, the BL-induced conformational changes in the LOV domain lead to the dissociation of the bZIP domain from the LOV surface (Fig. 6). We successfully visualized the loosening and elongation of the bZIP domain under BL illumination. In addition, we revealed that the elongated bZIP domain fluctuates around the LOV domain (Fig. 5). This direct visualization of such dynamic behavior of the domain structures in TFs has not previously been possible with any other technique and was achieved by HS-AFM for the first time in the present study. The fast diffusion of PZ monomers can increase collisions with other monomers (Fig. 3). In addition, the elongation and fluctuation of the bZIP domain can increase contact with other monomer because such a bZIP domain can have a greater capture radius for a specific binding site than the entangled state with restricted conformational freedom. This dynamic behavior of the bZIP domain is considered to contribute to the rapid formation of the stable dimer (Fig. 6). Previous studies on bZIP have proposed the fly-casting mechanism, where the fluctuation of a flexible bZIP domain accelerates the search for specific target DNA^42,43^. Our results suggest that this mechanism can also be extended to TF dimerization processes. Thus, PZ dimerization is accurately regulated by the autoinhibition due to the intramolecular interaction and is facilitated by the dynamic fluctuation of the flexible bZIP domain because of the BL-induced conformational change. Such autoinhibition due to intramolecular interactions has also been suggested for the other TFs^44,45^. Activating transcription factor 2 (ATF2)^46,47^ and CCAAT/enhancer binding protein-β (CEBPB)^48,49^ are examples of medically important TFs involved in various diseases such as cancer-cell multiplications and inflammatory processes. We expect HS-AFM to be useful for studying the unknown regulatory mechanism of such TFs through visualization of the domain structures and the corresponding single-molecular dynamics. Moreover, this technique is widely expected to be applicable for observing DNA-binding processes of TFs. In conclusion, we have revealed the significant role of the bZIP domain in the dimerization process for the photoresponsive TF photozipper. Our results demonstrate the enormous potential of HS-AFM imaging for transcriptional studies.

## Methods

### Sample preparation

Recombinant wild-type PZ and mutant S_2_C proteins were prepared as described previously^16^. In brief, *Escherichia coli* cells expressing recombinant PZ proteins were harvested by centrifugation and disrupted by sonication. After the cell debris was removed by centrifugation, recombinant proteins were purified twice by binding to and eluting from a Ni Sepharose 6 Fast Flow column (GE Healthcare) and a HiTrap Heparin HP column (GE Healthcare) according to the manufacturer’s instructions. The recombinant proteins were stored at 4°C in a solution (Buffer-A) containing 400 mM NaCl, 20 mM Tris-HCl (pH 7.0), 1 mM dithiothreitol (DTT), and 0.2 mM phenylmethylsulfonyl fluoride (PMSF). The purity of the PZ recombinant proteins was confirmed by SDS-PAGE, and the concentrations of the recombinant proteins were determined from the absorbance at 447 nm using an extinction coefficient of 13,000 M^−1^ cm^−1 16^. Samples for HS-AFM observations were diluted in a solution (Buffer-B) containing 1 mM NaCl, 20 mM Tris-HCl (pH 7), 1 mM DTT, and 0.2 mM PMSF.

### HS-AFM observation of PZ molecules

Imaging was performed with a laboratory-built HS-AFM apparatus, which is an improved version of the previously reported AFM^50^. The HS-AFM apparatus was installed in a darkroom. HS-AFM images were acquired using tapping mode. To detect cantilever deflection, we used an optical beam deflection detector equipped with an infrared laser (980 nm) to avoid exciting the PZ. The laser beam was focused onto a small cantilever [0.1 ≤ *k* ≤ 0.2 N/m, 800 ≤ *f* ≤ 1200 kHz in solution (Olympus)] using a 40× objective lens. The AFM styli were placed on each cantilever by electron-beam deposition and were sharpened by plasma etching. A 1.5 mm-diameter mica disk was glued onto a sample stage made of quartz glass. A sample droplet with a volume of 1.5 μL was placed on the freshly cleaved mica surface and incubated for 3 min. They were then rinsed with Buffer-B solution. These preparations for HS-AFM imaging were conducted under dim red light for the experiments under dark conditions and under blue LED light (450 nm) for those under light conditions. HS-AFM measurements were performed in Buffer-B solution at room temperature. For the photoexcitation of PZ samples during HS-AFM imaging, blue laser (450 nm) irradiation was performed through a 40× objective lens. The intensity measured at the exit of the objective lens was approximately 2 mW.

### Data analysis of HS-AFM images

The molecular volume and the mass center position of each imaged PZ molecule were analyzed using the following procedures. First, a Gaussian filter was applied to the HS-AFM images to reduce noise. Second, the grayscale of the images was digitized using a threshold determined by the half-maximum height to detect the boundary of a PZ molecule. The pixels with grayscales greater than the threshold were defined as the PZ molecular region (region of interest; ROI). The molecular volume of a PZ molecule was calculated by integrating the volume of each pixel within the ROI. The mass center position of a PZ molecule was calculated using the *X, Y*, and *Z* coordinates of pixels in the ROI.

The contour length of the bZIP-linker region was analyzed using the following procedure. First, the ROI of a PZ monomer region was determined. Second, a skeleton line was calculated in the ROI so that the distance to the border was the same on both sides of the skeleton line^51^. Third, the height profiles along the skeleton line were displayed as shown in Supplementary Fig. 5. In this profile, the domain boundary in a PZ molecule were determined by the half-maximum height. We then defined the upper and lower side of the boundary as the LOV domain and the bZIP-linker region, respectively. Finally, the length of the skeleton line in the bZIP-linker region was analyzed as the contour length of the bZIP-linker region.

## Data Availability

The complete amino acid sequence of PZ was deposited in the DDBJ database (accession number LC629158).

## Acknowledgements

This work was supported by JSPS KAKENHI Grant Numbers 20H03223 (to H.Y.) and 19K06586 (to O.H.), and 19H05789, 21H01812, 21K18876 (to M.A.), PRESTO, JST JPMJPR15FD, Multidisciplinary Research Laboratory System of Osaka University, the Osaka University Program for the Support of Networking among Present and Future Researchers, the joint research program of Biosignal Research Center, Kobe University 192005, and Takeda Science Foundation (to H.Y.).

## Author contributions

H.Y. and O.H. conceptualized the study. A.T. and H.Y. designed the experiments. O.H prepared the protein samples. A.T. and H.Y. performed and analyzed the HS-AFM experiments. H.Y., O.H. and M.A. administrated and supervised the study. A.T., H.Y. and O.H. wrote the paper.

## Ethics declarations

Competing interests

The authors declare no competing interests.

## Notes

### Competing Interest Statement

The authors have declared no competing interest.

